# Conserved and transcript-specific functions of the RESC factors, RESC13 and RESC14, in kinetoplastid RNA editing

**DOI:** 10.1101/2022.07.27.501750

**Authors:** Katherine Sortino, Brianna L. Tylec, Runpu Chen, Yijun Sun, Laurie K. Read

## Abstract

Uridine insertion/deletion RNA editing is an extensive post-transcriptional modification of mitochondrial mRNAs in kinetoplastid organisms, including *Trypanosoma brucei*. This process is carried out using *trans*-acting gRNAs and complex protein machinery. The essential RNA Editing Substrate Binding Complex (RESC) serves as the scaffold that modulates protein and RNA interactions during editing, and contains the Guide RNA Binding Complex (GRBC), the RNA Editing Mediator Complexes (REMCs), and organizer proteins. Despite the importance of RESC in editing, the functions of each protein comprising this complex are not completely understood. Here, we further define the roles of a REMC protein, RESC13, and a RESC organizer, RESC14, using high-throughput sequencing on two large pan-edited mRNAs, A6 and COIII. When comparing our analyses to that of a previously published small pan-edited mRNA, RPS12, we find that RESC13 has conserved functions across the three transcripts with regards to editing initiation, gRNA utilization, gRNA exchange, and restricting the formation of long mis-edited junctions that likely arise from its ability to modulate RNA structure. However, RESC13 does have transcript-specific effects on the types of long junctions whose formation it restricts. RESC14 has a conserved effect on gRNA utilization across the three transcripts analyzed, but has transcript-specific effects on editing initiation, gRNA exchange, and junction formation. Our data suggest that transcript-specific effects of both proteins are due to differences in transcript length and sequences as well as transcript-specific protein interactions. These findings highlight the importance of studying multiple transcripts to determine the function of editing factors.

## INTRODUCTION

The kinetoplastid parasite, *Trypanosoma brucei*, is the etiologic agent of Human African Trypanosomiasis (Bilbe 2015; Franco et al. 2020). Kinetoplastids are named for their unique mitochondrial DNA, called the kinetoplast or kDNA, which contains thousands of ∼1kb minicircles catenated with a few dozen ∼23kb maxicircles (Jensen and Englund 2012). Maxicircles encode rRNAs and subunits of respiratory complexes; however, 12 of the 18 protein-coding genes are considered cryptogenes because they do not encode functional reading frames (Aphasizheva et al. 2020). These mRNAs undergo an extensive posttranscriptional modification in which uridines (Us) are precisely inserted, and less frequently deleted, by a process called U insertion/deletion (U-indel) RNA editing (Cruz-Reyes et al. 2018; Zimmer et al. 2018; Aphasizheva et al. 2020). This process is essential for the survival of both procyclic (PF) and bloodstream form *T. brucei*, and is conserved across the Kinetoplastea (Schnaufer et al. 2001; Worthey et al. 2003; Tarun et al. 2008; David et al. 2015). In *T. brucei*, three of the 12 edited mRNAs only need a few dozen or less U insertions/deletions, and these are known as moderately edited (Aphasizheva et al. 2020). In contrast, nine mRNAs are termed pan-edited as they require extensive editing throughout their lengths, entailing hundreds of U insertions.

The sequence information that directs U-indel editing is contained in small guide RNAs (gRNA) that are primarily encoded in kDNA minicircles (Blum et al. 1990; Seiwert and Stuart 1994). Editing initiates when the anchor region of the first gRNA hybridizes to a portion of the 3’ never-edited region of the mRNA, while the rest of the gRNA forms an imperfect duplex with the mRNA. The editing machinery modifies the mRNA based on the gRNA’s coding region until there is complete gRNA-mRNA complementarity through Watson-Crick and G:U basepairing. The first gRNA is then removed by an unknown mechanism, and the next gRNA forms an anchor duplex with the newly edited mRNA sequence (Maslov and Simpson 1992). This process continues in the general 3’ to 5’ direction until the mRNA is fully edited. U-indel RNA editing is considered to be relatively inefficient because the bulk of steady-state mitochondrial mRNAs are partially edited, with only a very small number of fully edited mRNAs detectable (Koslowsky et al. 1991; Simpson et al. 2016; Gerasimov et al. 2018; Kumar et al. 2020; Smith et al. 2020; Dubey et al. 2021). Complicating matters, greater than 90% of these partially edited mRNAs contain regions of mis-editing termed junctions at the leading edge of a correctly sequence. Junctions are edited sequences of variable length that do not match the canonical, fully edited sequence and are thought to represent both areas of active editing that will be re-edited to the correct sequence and dead-end products that likely arise by non-cognate gRNA usage (Koslowsky et al. 1991; Simpson et al. 2016; Zimmer et al. 2018).

U-indel RNA editing is carried out by a holoenzyme complex composed of three dynamically interacting complexes: RNA Editing Core Complexes (RECCs), RNA Editing Substrate Binding Complex (RESC), and RNA Editing Helicase 2 Complex (REH2C) (Aphasizheva et al. 2020). Three related RECCs contain the enzymes needed to catalyze endonuclease cleavage, U insertion/deletion, and RNA ligation (Carnes et al. 2011; McDermott et al. 2016). However, purified RECC lacks associated RNA and is very poorly active *in vitro* (Rusche et al. 1997; Carnes et al. 2012). The RECCs interact transiently with REH2C and RESC through RNA interactions (Aphasizheva et al. 2014; Kumar et al. 2016; Aphasizheva et al. 2020). REH2C contains a DEAH/RHA type RNA helicase and two cofactors, and facilitates substrate-specific and site-preferential changes in total editing on transcripts associated with RESC (Kumar et al. 2016; Kumar et al. 2020). RESC contains ∼20 proteins arranged into modules: the Guide RNA Binding Complex (GRBC), the heterogeneous RNA Editing Mediator Complexes (REMCs), and in some models, the Polyadenylation Mediator complex (PAMC) (Aphasizheva et al. 2020; Dubey et al. 2021). Its GRBC module contains seven proteins that interact in an RNA-independent manner and appears relatively stable and homogeneous (Ammerman et al. 2012; Aphasizheva et al. 2014). In contrast, the REMCs represent a heterogenous, related set of complexes that interact with GRBC in an RNA-dependent manner (Aphasizheva et al. 2014; Simpson et al. 2017). In addition to these modules, RESC also contains a number of factors defined as RESC organizers: RESC8, RESC10 and RESC14. These proteins are not components of GRBC, REMCs or PAMC, but are important for modulating proper protein-protein and protein-RNA rearrangements during editing (McAdams et al. 2018; McAdams et al. 2019; Dubey et al. 2021). Importantly, purified REH2C and RESC include stably associated RNA (Weng et al. 2008; Madina et al. 2014). Thus, it is envisioned that RESC serves as the RNA editing scaffold, while the RECCs interact more transiently with RESC, REH2C, and the associated RNAs to catalyze editing (Aphasizheva et al. 2014; McAdams et al. 2018; Dubey et al. 2021).

Despite the critical importance of RESC in U-indel editing, the functions of its component proteins are incompletely understood. One reason for this is that the detailed impact of RESC factor knockdown on U-indel editing at the RNA sequence level has only been analyzed on a handful of small or moderately edited mRNAs and domains (Simpson et al. 2017; McAdams et al. 2018; McAdams et al. 2019; Tylec et al. 2019; Dubey et al. 2021). Thus, it is not known if the functions identified on these small mRNAs are conserved, or if some RESC factors, like REH2C (Kumar et al. 2020), have transcript-specific functions. In this study, we aim to further understand the role of two distinct and essential RESC proteins, RESC13 (formerly TbRGG2; Tb927.10.10830) and RESC14 (formerly MRB7260; Tb927.9.7260), in U-indel RNA editing using high-throughput sequencing (HTS) analysis of larger pan-edited mRNAs. RESC13 is an abundant REMC protein that strongly interacts with proteins of both REMC and GRBC (Ammerman et al. 2012; Aphasizheva et al. 2014; Simpson et al. 2017). RESC13 binds RNA as a dimer through its RRM domain and modulates RNA-RNA structure through annealing and melting activities, which are essential for its role in 3’ to 5’ editing progression (Fisk et al. 2008; Ammerman et al. 2010; Foda et al. 2012; Simpson et al. 2017; Travis et al. 2019). RESC14 is a RESC organizer that is particularly interesting because it does not bind RNA and has RNA-inhibited interactions with some RESC factors, suggesting it may compete with RNA for protein binding (McAdams et al. 2018). In this study, we employ HTS along with bioinformatic analysis using the Trypanosome RNA Editing Alignment Tool (TREAT) (Simpson et al. 2016; Simpson et al. 2017) on two long pan-edited mRNAs: ATPase synthase 6 (A6 pre-edited: 401 nucleotides (nt), fully edited: 820 nt) and cytochrome oxidase subunit III (COIII pre-edited: 463 nt, fully edited: 969 nt). We then compare their analyses to new and previously published analyses of the small pan-edited mRNA, ribosomal protein S12 (RPS12 pre-edited: 221 nt, fully edited: 325 nt) (Simpson et al. 2017; McAdams et al. 2018). These studies illuminate the roles of RESC13 and RESC14 during editing, and identify both conserved and transcript-specific impacts of these differentially functioning RESC proteins.

## RESULTS

### RESC13 does not function in editing initiation, while RESC14 has transcript-specific effects on editing initiation

We first asked whether RESC13 or RESC14 have transcript-specific effects on editing initiation, which manifest as an increase in pre-edited mRNAs after editing factor depletion. Previously, we showed by both qRT-PCR and HTS that there is no change in the level of pre-edited RPS12 mRNA after either RESC13 or RESC14 RNAi, indicating these proteins are not involved in editing initiation on RPS12 mRNA in PF cells (Fisk et al. 2008; Simpson et al. 2017; McAdams et al. 2018). To similarly analyze A6 and COIII mRNAs, we performed qRT-PCR analysis using primer sets specific to pre-edited and edited A6 and COIII mRNAs for two replicates each of RESC13- and RESC14-replete and depleted samples, after a two-day and three-day induction, respectively. (**Figs. 1A and 1B**). Our data confirm previously reported results showing that while RNAi of both factors decreases the levels of edited A6 and COIII mRNAs, the only observed increase in pre-edited mRNA is an approximately 4-fold increase in pre-edited COIII mRNA following RESC14 depletion (**Figs. 1A and 1B**) (Fisk et al. 2008; Simpson et al. 2017; McAdams et al. 2018). We also measured the total levels of A6 and COIII mRNAs, regardless of their editing status, revealing no changes in either mRNA in either cell line. This finding further supports that the observed increase in COIII pre-edited mRNA in RESC14 knockdowns is due to a change in the proportion of the mRNA population that has initiated editing and not to a change in overall mRNA abundance (**Figs. 1A and 1B**). These data suggest that RESC13 does not function in editing initiation, while RESC14 has transcript-specific effects on editing initiation.

**Fig. 1:**
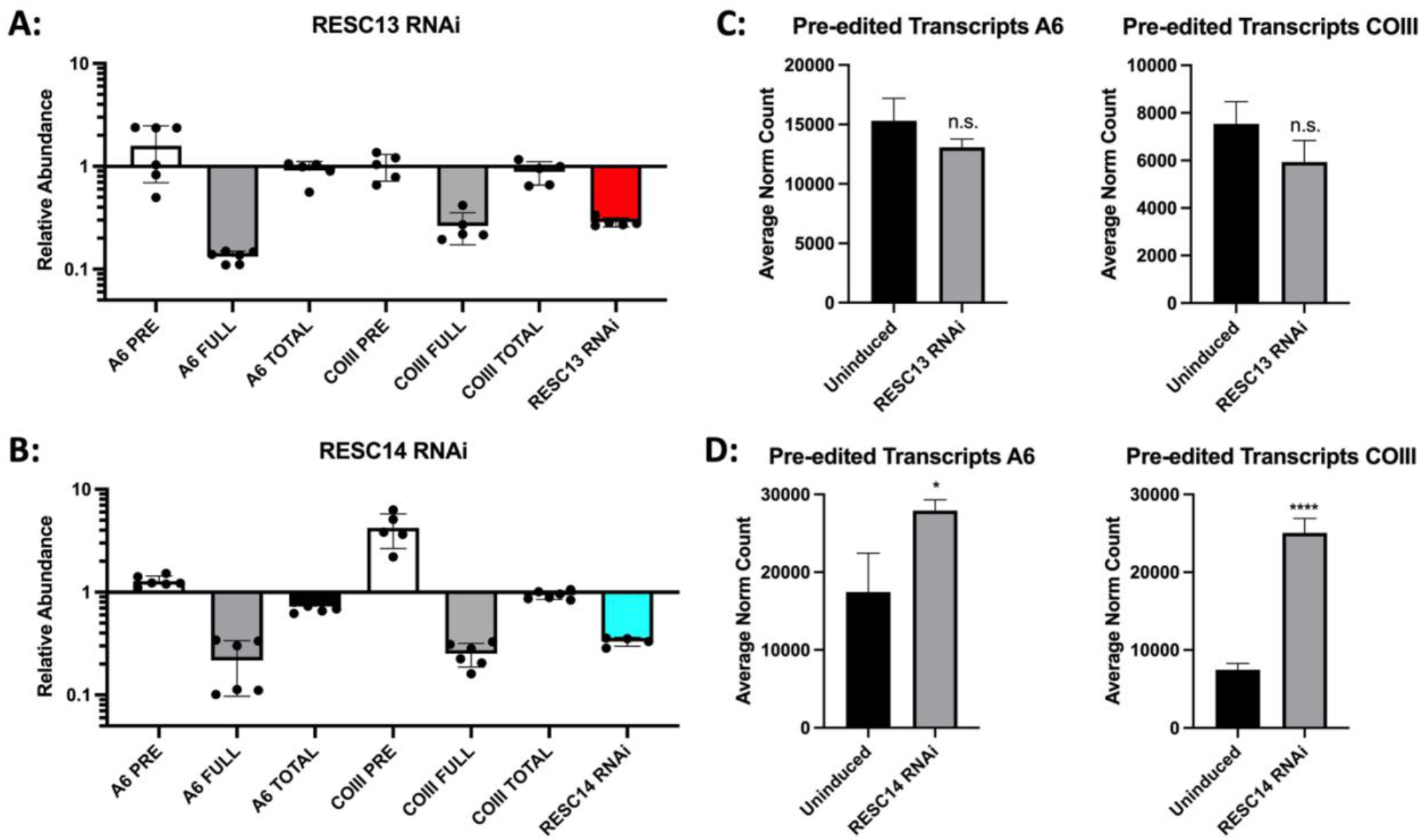
RESC13 is not involved in editing initiation, while RESC14 has transcript-specific effects in initiation. (**A**) qRT-PCR analysis of RNA isolated from uninduced and RESC13 RNAi-induced samples using primer sets designed to detect pre-edited, fully edited and total RNA levels of A6 and COIII. Relative abundance of each transcript in induced *vs*. uninduced RNAi cells is shown. Two biological replicates, each with three technical replicates, were performed. Replicate 1 RNA levels were normalized to 18S rRNA levels and replicate 2 levels were normalized to B-tubulin RNA levels. (**B**) Same as in **A**, but with RESC14 RNAi. RNA levels for both replicates were normalized to 18S rRNA levels. (**C**) The average number of normalized pre-edited A6 and COIII reads for uninduced samples and two RESC13 RNAi-induced samples determined through TREAT. (**D**) Same as in **C**, but with two RESC14 RNAi-induced samples.

To confirm the editing initiation phenotype determined through qRT-PCR, as well as to determine whether RESC13 and RESC14 have additional effects on editing progression on A6 and COIII mRNAs (see below), we turned to HTS. Paired-end Illumina MiSeq was used to sequence the 3’ region of pre-, partially- and fully-edited A6 and COIII from cells replete with and depleted of RESC13 or RESC14 (Simpson et al. 2016; Smith et al. 2020). The editing domains of pan-edited A6 and COIII are too large to be completely sequenced using MiSeq, so we amplified their 3’ end using a forward primer designed to a pre-edited sequence near the middle of the transcripts and a reverse primer designed to the 3’ never edited regions (Smith et al. 2020). “Fully” edited A6 refers here to mRNA intermediates that are canonically fully edited up to the forward pre-edited primer. Due to sequence ambiguities, “fully” edited COIII refers to transcripts matching canonical edited sequence up to ES 109, which is 5 ES 3’ of the forward primer (see Materials and Methods).

Libraries were obtained from two replicates each of induced RESC13 and RESC14 RNAi samples, and compared to two uninduced RESC14 or RESC13 RNAi cells, respectively, as well as five PF 29-13 cells from another study (compared to seven uninduced samples total) (Smith et al. 2020). Sequences were normalized to 100,000 reads and aligned using the TREAT algorithm, as previously described (Simpson et al. 2016; Simpson et al. 2017). Analysis of pre-edited mRNA levels through HTS showed that approximately 16% of A6 mRNAs and 7.5% of COIII mRNAs are pre-edited in uninduced cells (**Figs. 1C and 1D**), which is similar to the 14% pre-edited reads previously reported for RPS12 mRNA (Simpson et al. 2016). Upon RESC13 RNAi, we observed no significant change in the number of pre-edited sequences, confirming that RESC13 does not impact the initiation of editing on these transcripts (**Fig. 1C**). When RESC14 was depleted, we observed a >3-fold increase in pre-edited COIII mRNAs, consistent with qRT-PCR results (**Fig. 1D**). We also observed a modest but significant increase in pre-edited A6 mRNA, slightly more than observed with qRT-PCR (**Fig. 1D**). Overall, we conclude that RESC13 does not play a role in editing initiation. In contrast, RESC14 has transcript-specific functions in editing initiation, with no effect on RPS12, a minimal effect on A6 mRNA, and a significant effect on COIII mRNA.

RESC14 may have transcript-specific effects with regards to editing initiation due to differences in RNA structure of the three transcripts. Our data show that effects of RESC14 on editing initiation are very different between RPS12 and COIII mRNAs, with A6 being more intermediate (McAdams et al. 2018). Pre-edited RPS12 mRNA is just 221 nt, pre-edited A6 is 401 nt, and pre-edited COIII is 463 nt. Longer mRNAs likely permit more complex mRNA secondary structure formation and differences in mRNA-gRNA structure formation, each of which could contribute to challenges for the editing machinery to bind the mRNA and catalyze editing. RESC14 is important for modulating different protein-protein and protein-RNA interactions during editing (McAdams et al. 2018). Thus, it is possible that the more complex RNA structures of A6 and especially COIII mRNAs, necessitate RESC14 help to properly organize macromolecular interactions that allow editing to initiate. We also note that RPS12 and A6 are the only transcripts whose editing is required in bloodstream form *T. brucei*. Thus, these RNAs may have evolved to allow more permissive initiation compared to COIII mRNA.

### Impacts of RESC13 and RESC14 on gRNA utilization and exchange

We next wanted to determine how RESC13 and RESC14 impact 3’ to 5’ editing progression on A6 and COIII mRNAs and to identify conserved phenotypes of RESC13 and RESC14 across multiple transcripts (Simpson et al. 2017; McAdams et al. 2018). TREAT allows us to analyze and compare partially edited sequences within populations and to quantify features of editing progression as follows (Simpson et al. 2017). To determine the extent of canonical editing on a given read, TREAT defines the editing stop site, which is the 5’ most Editing Site (ES) after contiguous canonical editing that matches the fully edited sequence correctly (terminology in **Table 1**) (Simpson et al. 2016). The region 5’ of an editing stop site can either be pre-edited or a junction. Junctions are areas of mis-editing, in which the sequences match neither the pre-edited nor fully edited sequence, and they are found in a majority of our partially edited A6, COIII and RPS12 mRNAs (**Table 1**) (Simpson et al. 2016; Smith et al. 2020). TREAT defines junctions as extending from the 3’ most ES that does not match the fully edited sequence to the 5’ most editing site that shows any editing modification. Analysis using the TREAT algorithm allows us to determine whether depletion of a given factor causes editing to pause within or at the ends of distinct gRNA-directed blocks and to analyze the lengths and sequences of junctions (Simpson et al. 2016; Simpson et al. 2017). Focusing on these parameters, we evaluated the effects of RESC14 and RESC13 knockdown on A6 and COIII mRNAs.

**Table 1:** Glossary of terms.

| **Table 1: Glossary of terms** |  |
| --- | --- |
| Term | Definition |
| Editing Site (ES) | Any editing site between two non-U nucleotides that has the potential to be edited at the RNA level. ES are number in the 3' to 5' direction, as that is the way editing progresses. |
| Editing Stop Site | The 5' most (final) ES that matches the canonical fully edited sequence correctly. Any sequence 3' of the editing stop site is canonically fully edited. |
| Exacerbated Pause Site (EPS) | An editing stop site that significantly increases ($p_{adj} < 0.05$) when comparing two different RNA populations. This is typically determined when comparing RNAi cells to uninduced cells. |
| Junction | Mis-edited sequences found after the editing stop site- between 3' fully edited and 5' pre-edited sequences. |
| Junction Length (JL) | The number of ES a junction region spans. JL of 0 indicates there is no junction sequence, rather a completely pre-edited sequence 5' of the editing stop site. |

We began by defining the ES at which canonical editing pauses significantly more frequently in RESC13 and RESC14 knockdowns compared to their uninduced counterparts,. We refer to these sites as Exacerbated Pause Sites (EPSs) (**Table 1**). We analyzed EPS positions on A6 and COIII mRNAs relative to their fully edited sequence and the positions of previously reported gRNAs (**Fig. 2, diamonds; Fig. S1**) (Koslowsky et al. 2014). Most EPSs following RESC13 and RESC14 depletion are found throughout the lengths of gRNA-directed blocks on A6 and COIII mRNAs, similar to the EPSs on RPS12 mRNA after depletion of these proteins (Simpson et al. 2017; McAdams et al. 2018). The occurrence of numerous EPS within gRNA-directed blocks shows that RESC13 and RESC14 are needed for gRNA utilization on all three transcripts. Utilization of the initiator gRNA of COIII appears especially difficult upon knockdown of either of these factors as we observe robust EPS formation at the majority of ES throughout the gRNA-1 directed region (**Fig. 2B; blue and red shaded diamonds**).

**Fig. 2:**
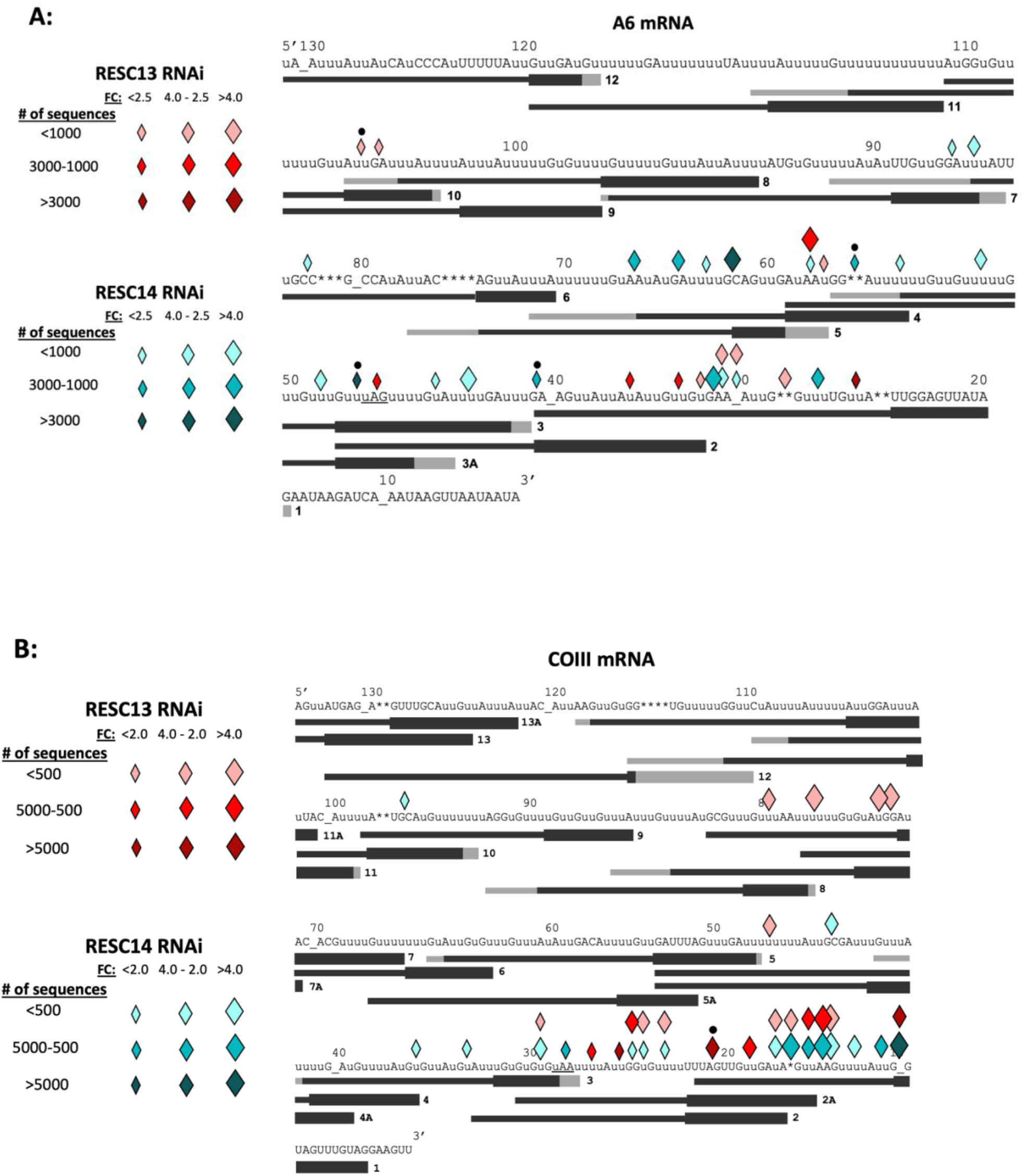
RESC13 and RESC14 functions in gRNA utilization and exchange. A6 (**A**) and COIII (**B**) edited mRNA sequences with the Exacerbated Pause Sites (EPS; diamonds) that arise upon RESC13 (red shades) and RESC14 (cyan shades) depletion. The average number of sequences found in the two induced replicates is represented by different diamond shading and the fold change of the induced samples to their uninduced controls are represented by different diamond sizes. The black dots above the diamonds represent EPS at gRNA ends. Black bars below the sequences represent cognate gRNAs as reported by Koslowsky et al. 2014, which are numbered in the 3’ to 5’ direction along the mRNA. gRNA anchors are depicted as the black bold lines where the grey regions denote range of variation of gRNA lengths across members of the same gRNA class. The uppercase Us are encoded in the mitochondrial genome, where the lowercase u’s represent inserted uridines during the editing process. Asterisks (*) denote sites where encoded uridines were deleted during the editing process. Underscores are shown for clarity in stretches of unedited sequence to align editing site numbers with the correct editing site. The stop codon is underlined.

In contrast to EPSs that occur within gRNA-directed blocks, EPSs at the ends of gRNA-directed blocks suggest effects of a given factor on gRNA exchange (Simpson et al. 2017). We previously reported that RESC13 and RESC14 RNAi led to EPSs at two (RESC13) and three (RESC14) gRNA ends on RPS12 mRNA (Simpson et al. 2017; McAdams et al. 2018). A role for RESC14 in gRNA exchange is also consistent with its presence in a small complex containing the gRNA binding proteins RESC1/2 (McAdams et al. 2018). To assess the potential impacts of RESC13 and RESC14 on gRNA exchange on A6 and COIII mRNAs, we determined whether any EPSs arising upon depletion of either of these factors were positioned at gRNA ends. RESC13 knockdown led to only one EPS at a gRNA end on either A6 or COIII mRNA (**Fig. 2; black circles above red diamonds**), suggesting little or no function in gRNA exchange for this factor on either mRNA. With regard to RESC14, its depletion led to EPSs at the ends of A6 gRNA-1, gRNA-2, and gRNA-3 (**Fig. 2A; black circles above blue diamonds**). EPSs towards the ends of A6 gRNA-4 and gRNA-6 could also be described as occurring at gRNA ends; however, we did not mark them as such because the extreme heterogeneity of the reported 3’ ends of these gRNA families (spanning three and five ESs, respectively; **Fig. 2A and B; grey line extensions on below gRNAs**) made this determination difficult. Thus, there may be more EPSs at gRNA ends in A6 mRNA upon RESC14 knockdown than we are highlighting. In contrast to its effect on A6 mRNA, RESC14 knockdown did not result in EPSs at gRNA ends on COIII mRNA, suggesting a transcript-specific phenotype of RESC14 in gRNA exchange. Little is known regarding the mechanisms of gRNA exchange, but again differences in intramolecular and mRNA-gRNA structures likely contribute to the transcript-specific differences observed here. Because RESC14 modulates macromolecular interactions during editing (McAdams et al. 2018), it is also plausible that it impacts transcript-specific editing accessory factors that function in gRNA exchange.

### RESC13 and RESC14 have both conserved and transcript-specific effects on junction formation

In addition to correct gRNA utilization and gRNA exchange, RESC factors can also affect the length and sequences of the mis-edited junctions that characterize the majority of partially edited mRNAs (Simpson et al. 2017; McAdams et al. 2018; McAdams et al. 2019; Dubey et al. 2021). A typical junction length is approximately 1-20 ES long, as that is the span of one gRNA. Junction lengths greater than 20 ES are considered long and likely arise through mRNA misfolding and/or extensive use of non-cognate gRNAs. A small fraction of partially edited sequences have no junctions, but rather transition directly from fully edited to pre-edited sequence. We characterize these sequences as having junction lengths of 0, and they represent a particular type of block in 3’ to 5’ editing progression. To further probe conserved and transcript-specific functions of RESC13 and RESC14 in editing progression, we asked whether the junctions that arise 5’ of editing stop sites in cells depleted of these factors are maintained across transcripts, initially focusing on junction length (**Table 1**). Using our HTS-TREAT libraries, we calculated the number of partially edited sequences that have junction lengths from 0, to 91 and above in the induced and uninduced RESC13 and RESC14 cells for A6, COIII and RPS12 mRNAs (**Fig. 3**). With regard to RPS12 mRNAs in RESC13 and RESC14 knockdown cells, we undertook new bioinformatic analyses of sequences that were previously reported (Simpson et al. 2017; McAdams et al. 2018), permitting comparison between this transcript and A6 and COIII mRNAs. Junction length analysis was performed previously on RPS12 mRNA in RESC13 RNAi cells; however, Simpson *et al*. only quantified short junction lengths of 0, 1, 2, 3, 4 and 5 across the entire RPS12 mRNA population (Simpson et al. 2017). Here, we quantify junction lengths 0 to >91, where ten editing sites are binned together (1-10, 11-20, etc.), allowing us to analyze more and longer junctions.

**Fig. 3:**
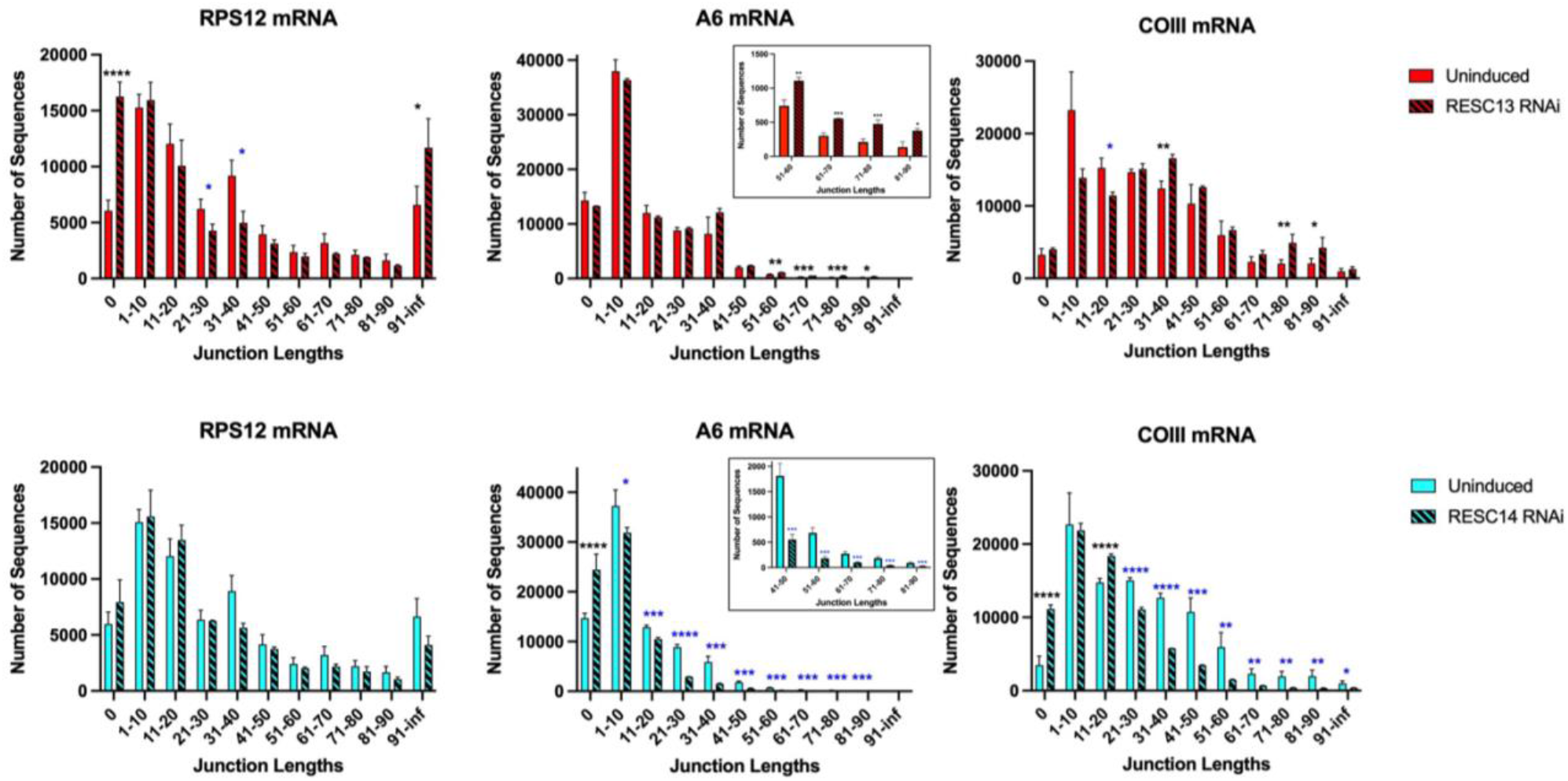
RESC13 and RESC14 have both conserved and transcript-specific effects on junction formation. The number of sequences for each junction length were quantified across the whole population of partially edited sequences for A6, COIII and RPS12 transcripts, in uninduced, RESC13 RNAi-induced (top; red) and RESC14 RNAi-induced (bottom; cyan) cells. Junction lengths are divided into 11 categories based on the number of editing sites spanned. Starting with length 0, then combining spans of 10 editing sites from lengths 1-90, followed by lengths 91 and above. Black asterisks denote junction lengths that were significantly increased in the RNAi samples relative to the uninduced, where blue asterisks denote junction lengths that were significantly decreased in the RNAi samples relative to the uninduced (padj<0.05). The insets for A6 mRNA show a close-up of the values for the longer junctions.

As previously described, RPS12 sequences with junction length 0 significantly increase across the entire populations upon RESC13 knockdown, indicating that RESC13 strongly promotes junction formation on this mRNA (**Fig. 3; top left**) (Simpson et al. 2017). A similar phenotype was also reported for the ND7-5’ mRNA following RESC13 knockdown (Simpson et al. 2017). In contrast, we observed no significant increase in junction length 0 sequences after RESC13 knockdown for A6 and COIII mRNAs (**Fig. 3; top middle and right**). We next determined the effects of RESC13 on different length junctions. We previously reported that junction lengths >50 significantly increased at specific EPSs on RPS12 mRNA after RESC13 RNAi (Simpson et al. 2017). Here, analyzing junctions across the entire RPS12 mRNA population, we show that the number of sequences with junction lengths >91 significantly increase after depleting RESC13 (**Fig. 3; top left**). From these data, we conclude that RESC13 plays another role in editing in addition to aiding in small junction formation, in this case restricting the formation of extremely long junctions on RPS12 mRNA. Strikingly, long junctions from lengths 51-90 (A6 mRNA) and 71-90 (COIII mRNA) also increase across A6 and COIII mRNA populations after RESC13 depletion, (**Fig. 3; top middle and left)**, highlighting a conserved function of RESC13 with regards to hindering the formation of long junctions. We next compared junction lengths across transcripts in RESC14 replete and depleted cells. We previously reported that RESC14 knockdown caused a small and statistically insignificant increase in the junction length 0 population of RPS12 transcripts, and a similarly small but insignificant decrease in junctions 50 ES and greater (McAdams et al. 2018). Our expanded analysis of RPS12 mRNA recapitulates these findings (**Fig. 3; bottom left**). In contrast, for A6 and COIII mRNAs, we observe that sequences with junction length 0 significantly increase with RESC14 RNAi, where all other sequence populations with junction lengths from 1-90 significantly decrease (**Fig. 3; bottom middle and right**).

Overall, A6 and COIII mRNAs behave more similarly with regard to junction lengths, while RPS12 mRNA is often an outlier. When examining the impact of RESC13 depletion, two different effects on junction formation were observed. The key role of RESC13 in restricting the formation of long junctions is conserved across all three transcripts. In contrast, RESC13’s role in facilitating short junctions was manifest only on RPS12 mRNA and absent in A6 and COIII mRNAs. RESC14 depletion substantially impacts junctions of all lengths in A6 and COIII mRNAs, indicating that RESC14 is especially critical for any 3’ to 5’ progression of editing on these mRNAs, including the mis-editing that characterizes junctions. This does not appear to be the case for RPS12 mRNA, where no significant effects on junction formation were observed upon RESC14 knockdown. However, we do note a similar, but statistically insignificant trend for RPS12 mRNA, where junction length 0 sequences slightly increase and long junctions slightly decrease. Overall, our data suggest that RESC14 is critical for any forward motion of editing, with junctions on longer transcripts being more sensitive than those on the shorter RPS12.

### RESC13 depletion leads to distinct types of long junctions on different mRNAs

Above, we showed that long junctions increase after RESC13 RNAi on all three mRNAs examined (**Fig. 3**). To understand what this means mechanistically, and to determine if long junctions form similarly on all three mRNAs, we determined the junction sequence characteristics of these mRNAs. To do so, we identified the editing stop sites at which junction lengths greater than 20 significantly increase after RESC13 RNAi on A6 (**Fig. 4A**), COIII and RPS12 (**Figs. S2A and S2B**) mRNAs. **Figs. 4A, S2A**, and **S2B** show the numbers of reads with junctions spanning 20 ES or more at each of these sites in RESC13 replete and depleted cells. Not all of these editing stop sites are EPSs (**Fig. 2**), although in each case over half are. Having identified editing stop sites with a significant increase in long junctions following RESC13 RNAi, we then analyzed the top 20 sequences at each of these sites to determine the sequence characteristics of the junctions and placed them into categories (**Fig. 4B-C**). Based on the TREAT algorithm, a junction extends from the 3’ most ES that does not match the fully edited sequence to the 5’ most editing site with any editing modification (**Table 1**). **Fig. 4B** illustrates several types of sequences that have been observed to contribute to the long junction phenotype.

**Fig. 4:**
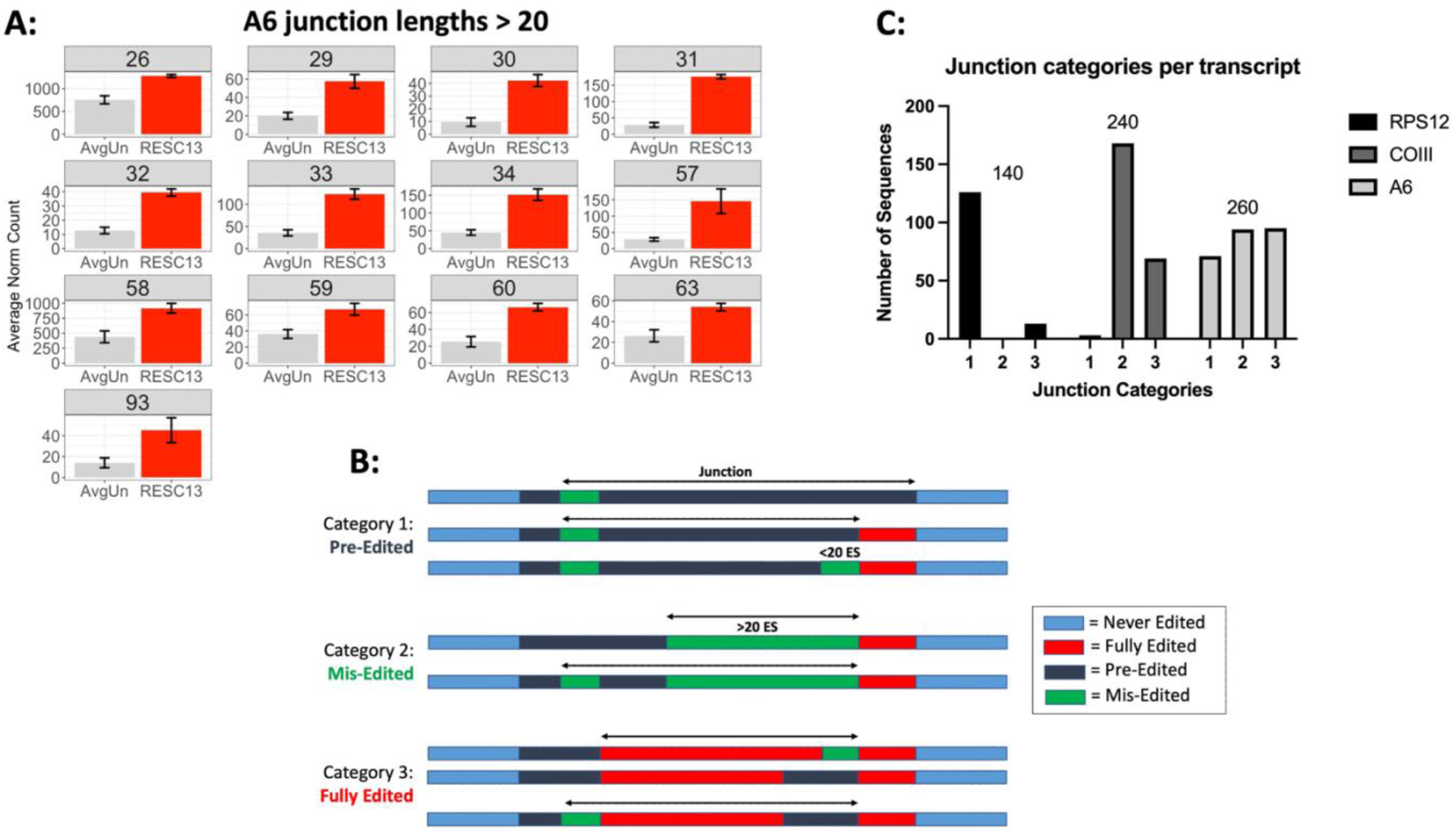
RESC13 hinders the formation of distinct classes of long junctions on different transcripts. (**A**) Numbers of reads at each A6 mRNA editing stop sites at which junctions with lengths greater than 20 ES increase upon RESC13 RNAi compared to uninduced cells. (**B**) Schematic of the different junction categories observed in RPS12, COIII, and A6 mRNA sequences that arise 5’ of the editing stop sites having JL>20 upon RESC13 RNAi. Three categories were identified: (1) mainly pre-edited, (2) mainly mis-edited, and (3) mainly fully edited. See key for clarification of the different colored bars. (**C**) The top 20 junction sequences that arise 5’ of editing stop sites where junctions greater than 20 ES long increase with RESC13 RNAi for A6, COIII and RPS12 mRNAs sorted into the three different categories. The numbers above the bars represent the total number of sequences analyzed for that transcript. Since there were a large number of JL>20 editing stop sites in COIII mRNA (Fig. S2A), only editing stop sites at which the normalized number of sequences in the induced cells was above 200 were analyzed.

As aforementioned, we previously showed that junction lengths >50 significantly increased at specific EPSs on RPS12 mRNA after RESC13 RNAi (Simpson et al. 2017). Sequence analysis showed that these long junctions displayed disjoined editing, in which there was mis-editing far 5’ in the transcript in the absence of contiguous 3’ editing, indicating that the region of active editing was not properly constrained (**Fig. 4B; Category 1**) (Simpson et al. 2017). This type of sequence could be generated through abnormal mRNA folding, allowing a gRNA to direct editing many ES 5’ of its normal target ES or by a mis-anchored gRNA directing editing in a region far 5’ of the ongoing normal editing progression. We show here that this type of junction, defined as Category 1 (**Fig. 4B**), is indeed the main type of long junction that increases with RESC13 RNAi on RPS12 mRNA (**Fig. 4C**).

We next wanted to know if this type of junction is increased after RESC13 depletion in other transcripts, to identify conserved phenotypes. When analyzing the sequences at each editing stop site where junction lengths greater than 20 increased upon RESC13 RNAi for A6 and COIII mRNAs, we observed two different sequence patterns emerge (**Fig. 4B; Category 2 and 3**). The sequences under Category 2 displayed very long stretches of mis-edited sequence in the position typical of a normal junction. Some of these sequences with long mis-edited regions had stretches of similar edited sequences at their 3’ ends that did not correspond to canonically, fully edited. These sequences were assumed to be alternative sequence, potentially directed by an alternative or non-cognate gRNA, followed by a true junction. Category 3 are sequences that are primarily fully edited, with either mis-edited or pre-edited sequence 3’ of a large stretch of fully edited sequence. Some of the sequences with a small amount of mis-editing 3’ could potentially be tolerated mistakes, either producing silent mutations or small conservating amino acid changes, and having enough correct editing that the next gRNA is still able to anchor (Simpson et al. 2016). Junctions with pre-edited sequence 3’ could represent gRNA misalignment or abnormal mRNA folding, allowing editing to be directed at a position 5’ of its canonical target sequence, again reflecting a lack of proper constraint of the region of active editing.

When characterizing the COIII mRNA long junctions, we found a majority of the sequences exhibited Category 2 (168 count), with barely any sequences falling under Category 1 (3 count) (**Fig. 4C**). Because long mis-edited sequences increase upon RESC13 RNAi, one interpretation is that RESC13 is important for hindering non-cognate gRNA usage on COIII mRNA. We also found a number of COIII junctions exhibiting Category 3 (69 count) (**Fig. 4C**), indicating RESC13 may also play a role in reducing the amount of tolerated editing mistakes on COIII mRNA. For A6 mRNA, observed approximately equal numbers of sequences exhibiting Category 2 (94 count) and Category 3 (95 count), while Category 1 had slightly fewer sequences (71 count) (**Fig. 4C**). This suggests that RESC13 is needed equally for constraining the region of active editing and hindering non-cognate gRNA utilization on A6 mRNA. Though RESC13 has transcript-specific effects on the types of long junctions that arise upon its depletion, these phenotypes may reflect similar RESC13 functions entailing modulation of RNA-RNA structure and constraint of non-cognate gRNA utilization.

When considering transcript length as a reason for why RESC13 functions differently on long junction formation of the three transcripts, we note that longer mRNAs have more opportunities for non-cognate gRNAs to anneal. We do not know the propensity for non-cognate gRNAs to interact with each mRNA or how more complex intra-mRNA structures enable or hinder gRNA annealing. With this said, Category 2 mainly likely arises due to non-cognate gRNA usage, and this category is mainly found on the long mRNAs, A6 and COIII (**Fig. 4C**). RESC13 therefore may play more of a role in hindering non-cognate gRNAs usage on A6 and COIII, due to their large size. Categories 1 and 3 are mainly formed when the region of active editing is not constrained, and these categories are found on RPS12 and A6 mRNAs (**Fig. 4C**). We hypothesize that the secondary and/or tertiary structures of these mRNAs are such that gRNAs can anchor and direct editing 5’ of their normal positions. Other factors we considered when addressing the difference of RESC13 function on long junction formation were a potential difference in the number of gRNAs that direct editing on each transcript, as well as a difference in the overlap of redundant gRNAs on each transcript. However, RPS12, A6 and COIII have a comparable number of gRNAs that direct editing in the region of interest with similar levels of gRNA redundancy or overlap. Thus, these features cannot account for differential RESC13 function with regard to long junction phenotypes on these three transcripts.

## CONCLUDING REMARKS

Here, we show through HTS and bioinformatic analysis that transcript lengths and sequences affect the necessity for or function of distinct RESC factors at different steps of editing. This highlights the importance of studying multiple transcripts when studying editing factor function. In addition to highlighting the role of mRNA identity in RESC function, our studies also provided insight into the functions of two RESC factors, RESC13 and RESC14. With regard to RESC13, to date we have analyzed two moderately edited mRNAs, three pan-edited mRNAs and one small mRNA editing domain by HTS in PF RESC13 RNAi cells, and in no case did we observe any effect on editing initiation (Simpson et al. 2017; Tylec et al. 2019). Thus, we are confident that RESC13 does not function in editing initiation, but rather has a primary function in promoting the 3’ to 5’ progression of editing. When addressing progression, RESC13 has a conserved function in constraining the region of active editing and promoting the usage of cognate gRNAs and/or restraining non-cognate gRNA utilization. This can likely be explained by the known biochemical functions of this protein, which includes RNA binding, unwinding and annealing activities (Fisk et al. 2008; Ammerman et al. 2010; Foda et al. 2012; Travis et al. 2019). It appears that RESC13’s main function is to promote correct intra- and inter-molecular RNA interactions during editing, making its modulation of RNA-RNA interactions imperative for all transcripts.

RESC14 has very different properties than RESC13. While RESC14 does not bind RNA and may compete with RNA for protein binding, it is involved in modulating proper protein-protein and protein-RNA interactions (McAdams et al. 2018). We find transcript-specific effects of RESC14 on editing initiation and potential role in gRNA exchange, based on observing EPSs at gRNA ends. This suggests that different transcripts may need different RESC14-mediated protein-protein interactions at these stages of editing. For example, different transcripts may require different REMCs or accessory factors, such as helicases or MRP1/2, and RESC14 may facilitate the interactions of these factors with the editing holoenzyme. We also found that RESC14 is critical in promoting 3’ to 5’ editing progression of editing, whether that be correct or mis-editing on most, if not all transcripts examined, suggesting it controls interactions of essential RESC players. Overall, these studies shed light on the functions of a REMC protein and a RESC organizer, allowing us to further understand the mechanism of action of the U-indel editing machinery.

## MATERIALS AND METHODS

### Cell culture

The cell lines used in this paper were derived from the procyclic form (PF) *T. brucei* 29-13 of the Lister 427 strain and were grown in SDM79 media supplemented with 10% fetal bovine serum, 50 µg/mL hygromycin, and 15 µg/mL G418 (Wirtz et al. 1999; Pelletier and Read 2003). The RESC14 and RESC13 RNAi cell lines were previously published (Ammerman et al. 2012; McAdams et al. 2018), and were grown in selective media with 2.5 µg/mL phleomycin.

### RNA isolation and qRT-PCR

RESC14 and RESC13 RNAi cells were grown either uninduced or induced with 4 µg/mL doxycycline for three days and two days, respectively. These time points were chosen as the target RNA was decreased while cell growth was not affected. Cells were harvested and total RNA was extracted using TRIzol reagent (Ambion) and phenol:chloroform extraction. The RNA was DNAse-treated for 1.5 hrs with a DNA-free DNAse kit (Ambion) and was reverse-transcribed to cDNA using random hexamer primers and the iScript reverse transcription kit (BioRad). To detect the levels of pre-edited, fully edited, and total transcripts (A6 and COIII), qRT-PCR was performed using established primers (Carnes et al. 2005; Fisk et al. 2008). Our total RNA primers detect the largest pool of mRNA for a given transcript as the forward primer is designed to the 5’ never-edited region and the reverse primer is designed to a far 5’ pre-edited sequence of a specific mRNA (McAdams et al. 2018). These primers will amplify all pre-edited and a majority of partially edited mRNAs but will not amplify fully edited. Since a small portion of mRNAs are edited extremely 5’ (Simpson et al. 2016; Simpson et al. 2017), these primer sets detect a majority of a specific mRNA population. Levels of RESC14 and RESC13 RNAi were detected using previously published RESC14- and RESC13-specific primers (Foda et al. 2012; McAdams et al. 2018). qRT-PCR results were analyzed using BioRad CFX Manager software and the RNA levels were normalized to levels of 18S or beta tubulin using the standard curve method. All results shown reflect two biological replicates with three technical replicates of the qRT-PCR reaction.

### Preparation of RNA for high-throughput sequencing

PF *T. brucei* RESC14 and RESC13 RNAi cell lines were grown in the presence or absence of 4 µg/mL doxycycline for three days and two days, respectively. Total RNA was extracted using Trizol, followed by phenol:chloroform extraction and DNAse treatment. Two biological replicates were performed for each cell line. DNAse treated RNA (1.2 µg) was used to generate cDNA with the Superscript III Reverse Transcriptase Kit (Invitrogen) and previously published gene specific primers to the 3’ region of the editing domain for cytochrome oxidase subunit III (COIII) and ATPase subunit 6 (A6) (Smith et al. 2020). Due to sequencing discrepancies at the very 5’ end of the COIII sequenced region, we analyzed COIII sequences up to 5 ES 3’ of the true 5’ end (ES 109); “fully edited” COIII refers to transcripts matching canonical edited sequence up to ES 109. To ensure the relative abundance of unique fragments was maintained, the linear range of the gene specific cDNA was determined through qRT-PCR. The amplicons for MiSeq sequencing were then generated using PCR amplification of the cDNA with the correct cycle number corresponding to the center of the linear range. The PCR amplicons were purified using the Illustra GFX PCR cleaning kit and eluted into 20 µl of 10mM Tris-HCl, pH.8. Library preparation and paired-end Illumnia Mi-Seq was used for high throughput sequencing of amplicons (see below). For A6 and COIII, two replicates of the induced RESC14 and RESC13 RNAi cell lines were used and compared to two uninduced RESC14 or RESC13 RNAi cells, respectively, as well as five PF 29-13 cells from another study (Smith et al. 2020). Sample preparation for ribosomal protein S12 (RPS12) was previously done and controls were described for RESC14 (McAdams et al. 2018) and RESC13 RNAi (Simpson et al. 2017).

### Library construction and sequencing

cDNA was quantified using the high sensitivity Qubit fluorescence assay (Invitrogen), and the sizes of products were confirmed using the Agilent Fragment Analyzer using standard sense reagents. A 50 µl index PCR reaction was carried out to attach dual indices and Illumina sequencing adapters. Twenty-five microliter of 2 X KAPA HiFi HotStart Ready mix was combined with 5 µl Nextera XT Index primer 1 (N7xx) and Index primer 2 (S5xx) and added to 2 ng of cDNA for the PCR reaction. AMPure XP beads (Beckman Coulter Genomics) were used to purify the final libraries. Libraries were then quantified using the Qubit assay and Library Quantification kit (Kapa Biosystems). The Agilent Fragment Analyzer was used to analyze the sizes of the cDNA libraries and confirm the presence of appropriate size ranges for both the edited and unedited transcript sizes as well as ligated sequencing adapters. The libraries were quantified using a Qubit fluorimeter (Thermo Fisher Scientific) and normalized to 10 nM based on the concentration and amplicon fragment size. The libraries were pooled and sequenced using Illumina MiSeq 300 cycle Paired End sequencing.

### Sequence alignment using Trypanosome RNA Editing Alignment Tool (TREAT)

TREAT is a multiple sequence alignment and visualization tool that was developed in our laboratory to analyze uridine insertion/deletion RNA editing (Simpson et al. 2016). TREAT consists of a command-line alignment algorithm and a web-based interface for searching, viewing, and analyzing sequence results. TREAT is written in Go and freely available under the GPLv3 license at http://github.com/ubccr/treat. TREAT v0.03 (Simpson et al. 2017) was used in this study. All reads were aligned to the published pre-edited and fully edited A6 and COIII mRNA sequences (Smith et al. 2020), with the modification of COIII mRNA as described above. The number of standard reads (sequences with no non-T mismatches) and non-standard reads (sequences with non-T mismatches) are listed in (**Table S1**). For each sample, the standard reads were normalized to 100,000. This allows the relative abundance of each sample to be scaled such that they can be compared via their normalized counts (Simpson et al. 2016). The non-standard reads were excluded from the analysis. The new sequencing data for the A6 and COIII, RESC13 and RESC14 RNAi samples has been deposited in the Sequence Read Archive under accession number PRJNA862535. Previously published sequencing data is under accession number PRJNA597932 for the A6 and COIII PF 29-13 samples, SRP109103 for the RPS12 RESC14 RNAi samples, and SRP097727 for the RPS12 RESC13 RNAi and RPS12 control samples.

### Determination of significant increase of pre-edited transcripts

The number of pre-edited transcripts found in each sample were determined by TREAT analysis (Simpson et al. 2016). For A6 and COIII, two RESC14 and RESC13 induced replicates were compared to their two uninduced counterparts, along with five PF 29-13 samples (Smith et al. 2020) (seven uninduced total). The averages and standard deviations were calculated for each induced and uninduced controls. Significance of changes to pre-edited transcript levels upon RNAi was determined using student’s *t-*tests. The effects of knockdown of a given protein were considered significant if both induced replicates were significantly increased compared to their uninduced controls.

### Determination of exacerbated pause sites (EPSs) and junction lengths/sequences

Determination of EPSs was described previously (Simpson et al. 2017). Briefly, editing stop sites were considered EPSs when the increase in sequence abundance at that editing stop site was statistically significant (padj<0.05) in both induced replicates compared to their uninduced controls (**Table S2 and S3**). EPSs were further characterized by the average number of sequences found in the two induced replicates (different diamond shading) as well as the fold change compared to the average of their uninduced controls (different diamond sizes). Calculating the significance of EPS overlap between the RNAi of different proteins was previously described (Simpson et al. 2017). For analysis of junction lengths, the total number of sequences containing a junction length of 0, 1-10, 11-20, 21-30, 31-40, 41-50, 61-70, 81-90 and greater than 91 ES long across the entire A6 and COIII RNA populations, was determined. The average number of sequences for each induced and uninduced samples were calculated and plotted in R. To determine RPS12, A6, and COIII mRNA editing stop sites at which sequences with junction lengths greater than 20 ES long were significantly increased in the RESC13 RNAi samples compared to the uninduced controls, student’s *t-*tests in R was performed. The top 20 sequences with junction lengths 20 ES long or greater at these significantly increased ES were analyzed, and the junction sequences were characterized into one of three categories: 1) predominantly pre-edited sequence, 2) predominantly mis-edited sequence, and 3) predominantly fully-edited sequence.

## SUPPLEMENTAL DATA

**Fig. S1** reports the Exacerbated Pause Sites (EPS) on A6 and COIII after RESC13 and RESC14 RNAi. **Fig. S2** shows the sites where sequences with JL>20 increase after RESC13 RNAi on COIII and RPS12. **Table S1** shows the total fragments and unique sequences in the partially edited A6 and COIII sequence libraries. **Tables S2 and S3** report the determination of EPSs on A6 and COIII after RESC13 and RESC14 RNAi.

## ACKNOWLEDGEMENTS

We thank Dr. Joseph T. Smith for critical reading of the manuscript. We are also grateful to the University at Buffalo Center for Computational Research (especially Andrew Bruno) and the University at Buffalo Genomics and Bioinformatics Core (especially Jonathan Bard) for their support. This work was supported by NIH grants R01 GM129041 (to LKR) and R01 CA241123 (to YS).

